# Language evolution to revolution: the leap from rich-vocabulary non-recursive communication system to recursive language 70,000 years ago was associated with acquisition of a novel component of imagination, called Prefrontal Synthesis, enabled by a mutation that slowed down the prefrontal cortex maturation simultaneously in two or more children – the Romulus and Remus hypothesis

**DOI:** 10.1101/166520

**Authors:** Andrey Vyshedskiy

## Abstract

There is an overwhelming archeological and genetic evidence that modern speech apparatus was acquired by hominins by 600,000 years ago^1^. On the other hand, artifacts signifying modern imagination, such as (1) composite figurative arts, (2) bone needles with an eye, (3) construction of dwellings, and (4) elaborate burials arose not earlier than 70,000 years ago^2^. It remains unclear (1) why there was a long gap between acquisition of modern speech apparatus and modern imagination, (2) what triggered the acquisition of modern imagination 70,000 years ago, and (3) what role language might have played in this process. Our research into evolutionary origin of modern imagination has been driven by the observation of a temporal limit for the development of a particular component of imagination. Modern children not exposed to recursive language in early childhood never acquire the type of active constructive imagination called Prefrontal Synthesis (PFS). Unlike vocabulary and grammar acquisition, which can be learned throughout one’s lifetime, there is a strong critical period for the development of PFS and individuals not exposed to recursive language in early childhood can never acquire PFS as adults. Their language will always lack understanding of spatial prepositions and recursion that depend on the PFS ability. In a similar manner, early hominins would not have been able to learn recursive language as adults and, therefore, would not be able to teach recursive language to their children. Thus, the existence of a strong critical period for PFS acquisition creates an evolutionary barrier for behavioral modernity. An evolutionary mathematical model suggests that a synergistic confluence of three events (1) a genetic mutation that extended the critical period by slowing down the prefrontal cortex development simultaneously in two or more children, (2) invention of recursive elements of language, such as spatial prepositions, by these children and (3) their dialogic communications using these recursive elements, resulted in concurrent conversion of a non-recursive communication system of their parents to recursive language and acquisition of PFS around 70,000 years ago.

## Introduction

Association of Wernicke’s and Broca’s areas with language is well-known. Less common is realization that understanding of full recursive language depends on the lateral prefrontal cortex (LPFC). Wernicke’s area primarily links words with objects^3^, Broca’s area interprets the grammar and assigns words in a sentence to a grammatical group such as noun, verb, or preposition^3^, but only the LPFC can combine objects from memory into a novel mental image according to a sentence’s description^4,5^. The latter function is commonly called imagination. The term “imagination,” however, is ambiguous as it is regularly used to describe any experience that was generated internally. For example, dreaming is often described as an imaginary experience. However, dreaming has a completely different neurological mechanism, as dreaming is not controlled by the LPFC^6–8^. LPFC is inactive during sleep^6,8^ and patients whose LPFC is damaged do not notice change in their dreams^7^. In order to distinguish the imagination during dreaming from the conscious purposeful active LPFC-driven synthesis of *novel* mental images, we define the latter process as *Prefrontal Synthesis* (PFS, a.k.a. mental synthesis)^9,10^. PFS is completely dependent on an intact LPFC^11–16^ and patients with damage to LPFC often lose their PFS function confirming neurological dissociation between the two types of imaginary experience^2^.

Patients with damage to the LPFC and spared Broca’s area often present with a specific PFS deficit, that affects both their language and imagination. Joaquin Fuster calls their alteration in language “prefrontal aphasia”^15^ and explains that “although the pronunciation of words and sentences remains intact, language is impoverished and shows an apparent diminution of the capacity to ‘propositionize.’ The length and complexity of sentences are reduced. There is a dearth of dependent clauses and, more generally, an underutilization of what Chomsky characterizes as the potential for recursiveness of language”^15^. Alexander Luria calls this condition “frontal dynamic aphasia”^17^ and reports that as long as a conversation does not involve combination of objects, these patients look unremarkable. They do not lose their vocabulary and can keep a conversation going: “…patients with this type of lesion have no difficulty articulating words. They are also able to retain their ability to hear and understand most spoken language. Their ability to use numerical symbols and many different kinds of abstract concepts also remains undamaged… these patients had no difficulty grasping the meaning of complex ideas such as ‘causation,’ ‘development,’ or ‘cooperation.’ They were also able to hold abstract conversations. … They can repeat and understand sentences that simply communicate events by creating a sequence of verbal images”^18^. Luria further explains that their disability shows only when patients have to imagine several objects or persons in a *novel* combination (revealing the problem of PFS): “But difficulties developed when they were presented with complex grammatical constructions which coded logical relations. … Such patients find it almost impossible to understand phrases and words which denote relative position and cannot carry out a simple instruction like ‘draw a triangle above a circle.’ This difficulty goes beyond parts of speech that code spatial relations. Phrases like ‘Sonya is lighter than Natasha’ also prove troublesome for these patients, as do temporal relations like ‘spring is before summer’ [AV: space is commonly used to represent time and therefore PFS disability usually results in inability to understand temporal relationship as well]. …Their particular kind of aphasia becomes apparent only when they have to operate with groups or arrangements of elements. If these patients are asked, ‘Point to the pencil with the key drawn on it’ or ‘Where is my sister’s friend?’ they do not understand what is being said. As one patient put it, ‘I know where there is a sister and a friend, but I don’t know who belongs to whom’”^18^. In our research, we have found that simplest relational inquiries can often quickly elucidate PFS disability. Questions, such as “If a cat ate a dog, who is alive?” and “Imagine the *blue* cup inside the *yellow* cup, which cup is on top? can be consistently answered by four-year-old children but commonly failed by individuals with PFS disability^19^.

Crucially, PFS disability is not just a receptive language disorder, but a deficit in imagination that can be confirmed by nonverbal IQ tests. Individuals with PFS disability may have normal full-scale IQ, but commonly exhibit a selective and catastrophic deficit in tasks relying on PFS, e.g., matrix reasoning tasks requiring integration of multiple objects^12^, such as those shown in Figure 1. Persons with PFS disability invariably fail these integration tasks and, therefore, typically perform below the score of 86 in non-verbal IQ tests^20^.

**Figure 1.**
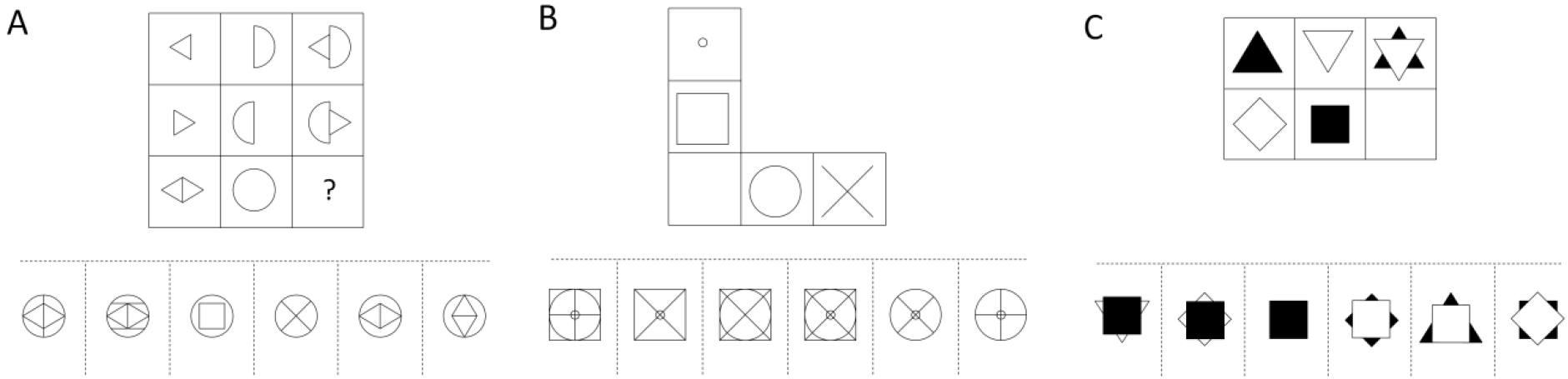
PFS disability goes beyond problems with understanding recursive language. This is the disability of one of the mechanisms of active imagination. Nonverbal tasks requiring imagining a novel combination of two or more objects is impossible in this condition. Typical IQ test tasks involving PFS of several objects: (A) requires the combination of two objects. The top two rows of the matrix indicate the rule: “the object in the right column is the result of the combination of the two objects shown in the left and middle row” (the solution in the 5^th^ square). (B) shows a question that relies on the PFS of four objects. (C) shows a question in which PFS of two objects has to be conducted according to the following rule specified in the top row: “the object in the middle column goes on top of the object in the left column” (the solution in the second square). Note that patients with PFC disability commonly have no problem with simpler performance IQ tasks, such as integration of modifiers^5^.

A deficit in imagination in otherwise unremarkable individuals is extremely counterintuitive for several reasons. First, most scientists have never met anyone with PFS disability as these individuals do not frequent university campuses and other privileged institutions. Second reason is that PFS was traditionally grouped with the other components of imagination, such as dreaming, spontaneous insight, and integration of modifiers, making it difficult to discuss the exact nature of a patient’s deficit, whose dreaming, spontaneous insight, and integration of modifiers remain normal. Finally, even when we see individuals with PFS disability, we tend to anthropomorphize, assume their normal PFS, and brush off their shortfall as a linguistic problem, memory deficit, or inattentiveness. However, when considered together with nonverbal IQ test, specifically those items that require integration of objects, there can be no mistake: as many as 18% of modern individuals exhibit PFS disability^5^. These individuals include some patients with damage to their PFC, low-functioning individuals with congenital neurological problems, as well as individuals with no congenital abnormality, who were not exposed to recursive language in early childhood (see below).

**Table 1.** Definitions used in the article

| Term | Definition |
| --- | --- |
| Prefrontal Synthesis (PFS) | PFS is the type of imagination that involves conscious, purposeful process of synthesizing novel mental images from two or more objects stored in memory |
| Active imagination | Active imagination includes several neurologically distinct components, such as PFS, Prefrontal Analysis, integration of modifiers and mental rotation. Active imagination completely depends on the LPFC. Active imagination is contrasted to spontaneous imagination, that includes dreaming, amodal completion, and spontaneous insight and does not depend on the LPFC. |
| Recursive language | Recursive language relies on a listener's PFS ability, to communicate infinite number of novel images with high fidelity. The hallmarks of recursive languages are spatial prepositions, verb tenses, nesting, and other recursive elements that facilitate language ability to flexibly describe various combinations of objects. |
| Non-recursive communication system | Before acquisition of PFS, our ancestors could not mentally arrange objects into novel combinations and therefore their communication system could not include spatial prepositions or recursion. While their non-recursive communication system probably had many words, it was essentially finite as it could not communicate as many novel images as a recursive language. |

### Comparison of Prefrontal Synthesis to linguistically defined Merge

PFS is defined neurobiologically as the conscious, purposeful process of synthesizing *novel* mental images from two or more objects stored in memory. There is no linguistically defined process that is neurobiologically equivalent to this distinct mechanism of imagination. The closest in spirit is Chomskyan Merge, defined as a process of combining any two syntactic objects to create a new one^21^. PFS, however, is different both in scope and underlying neurobiology. An individual does not need to know the names of objects in order to combine them mentally into a novel hybrid object or scene. One can mentally combine objects of strange geometrical shape that do not have names in any language. Merging of objects in mental space does not directly depend on knowledge of any language.

Even when language is used to direct PFS in the mind of interlocutor, PFS definition is different from Merge. For example, combination of an adjective and a noun is a Merge operation, but does not fall under PFS that must always involve combination of two or more independent objects. Furthermore, PFS, but not Merge, requires creating a *novel* mental hybrid object or scene. For example, a sentence ‘ship sinks’ can be understood by *remembering* a previously seen picture of a sinking ship and thus, completely avoiding the PFS process. Under Chomskyan theory, ‘ship sinks’, however, is considered a Merge operation since the sentence merges a determiner phrase ‘ship’ and a verb ‘sinks’ to create a sentence ‘ship sinks.’

In neurobiological terms, Merge operation is defined in such a way that it utilizes all three brain language regions: Wernicke’s area that primarily links words with objects; Broca’s area that interprets the grammar and assigns words in a sentence to a grammatical group such as noun, verb, or preposition^3^; and the LPFC that synthesizes the objects from memory into a novel mental image according to sentence’s description^4,5^. Crucially, PFS definition leaves out interpretation of grammar in the Broca’s area and leaves out linking words with objects in the Wernicke’s area. PFS definition limits it to the function of the LPFC. Thus, PFS is defined significantly more narrowly than the Merge operation in both neurological and linguistic terms.

The difference between PFS and Merge can be also highlighted by the process of learning a new language in adulthood: when one studies German, Spanish or Italian, one learns new words (Wernicke’s area) and new grammar rules (Broca’s area), PFS, however, does not change a bit. The same PFS abilities can be used to understand German, Spanish, or Italian sentences. PFS, in other words, is universal across all modern human languages, despite different words and grammar rules.

### Prefrontal Synthesis ability is essential for recursive language

All human languages *allow high fidelity transmission of infinite number of novel* images with the use of a *finite* number of words (here and later, words are understood as units of meaning, called sememes by linguists). The magic of using a finite number of words to communicate an infinite number of images completely depends on interlocutor’s ability to conduct PFS. When we describe a novel image (“My house is the second one on the left, just across the road from the church”), we rely on the interlocutor to use PFS in order to visualize the novel image. When we tell stories, we are often describing things that the interlocutor has never seen before (“That creature has three heads, two tails, large green eyes, and can run faster than a cheetah”) and we rely on the interlocutor to imagine the story in their mind’s eye. As Steven Pinker put it, “the speaker has a thought, makes a sound, and counts on the listener to hear the sound and to recover that thought”^22^. The importance of the PFS ability is best realized when one is attempting to understand sentences describing combination of objects. Consider the two sentences: “A dog bit my friend” and “My friend bit a dog.” It is impossible to distinguish the difference in meaning using words or grammar alone, since both words and grammatical structure are identical in these two sentences. Understanding the difference in meaning and appreciating the misfortune of the 1^st^ sentence and the humor of the 2^nd^ sentence depends on interlocutor’s PFS ability. Only after the LPFC forms these two different images in front of the mind’s eye, are we able to understand the difference between these two sentences. Similarly, recursive statements with nested explanations (“a snake on the boulder to the left of the tall tree that is behind the hill”) force interlocutor to use PFS to combine objects (a snake, the boulder, the tree, and the hill) into a novel scene. For this reason, linguists refer to modern full human languages as recursive languages.

This ability of recursive language to communicate an infinite number of novel images with the use of a finite number of words depends on interlocutor’s PFS capacity and also is facilitated by spatial prepositions, verb tenses, nesting, and other common elements of grammar. Consider, for example, the exponential ability of spatial prepositions to increase the maximum number of distinct images that can be communicated with high fidelity (Figure 2). In a language with no spatial prepositions or other recursive elements, 1000 nouns can communicate 1000 images to a listener. Adding just one spatial preposition allows for the formation of three-word phrases (such as: ‘a bowl behind a cup’ or ‘a cup behind a bowl’) and increases the number of distinct images that can be communicated from 1000 to one million (1000×1×1000, Figure 2). Adding a second spatial preposition and allowing for five-word sentences of the form object-preposition-object-preposition-object (such as: ‘a cup on a plate behind a bowl’) increases the number of distinct images that can be communicated to four billion (1000×2×1000×2×1000). The addition of a third spatial preposition increases the number of distinct images to 27 trillion (1000×3×1000×3×1000×3×1000), and so on. A typical language with 1000 nouns and 100 spatial prepositions can theoretically communicate 1000^101 x 100^100 distinct images. This number is significantly greater than the total number of atoms in the universe. For all practical purposes, an infinite number of distinct images can be communicated by a recursive language with just 1000 words and a few prepositions.

**Figure 2.**
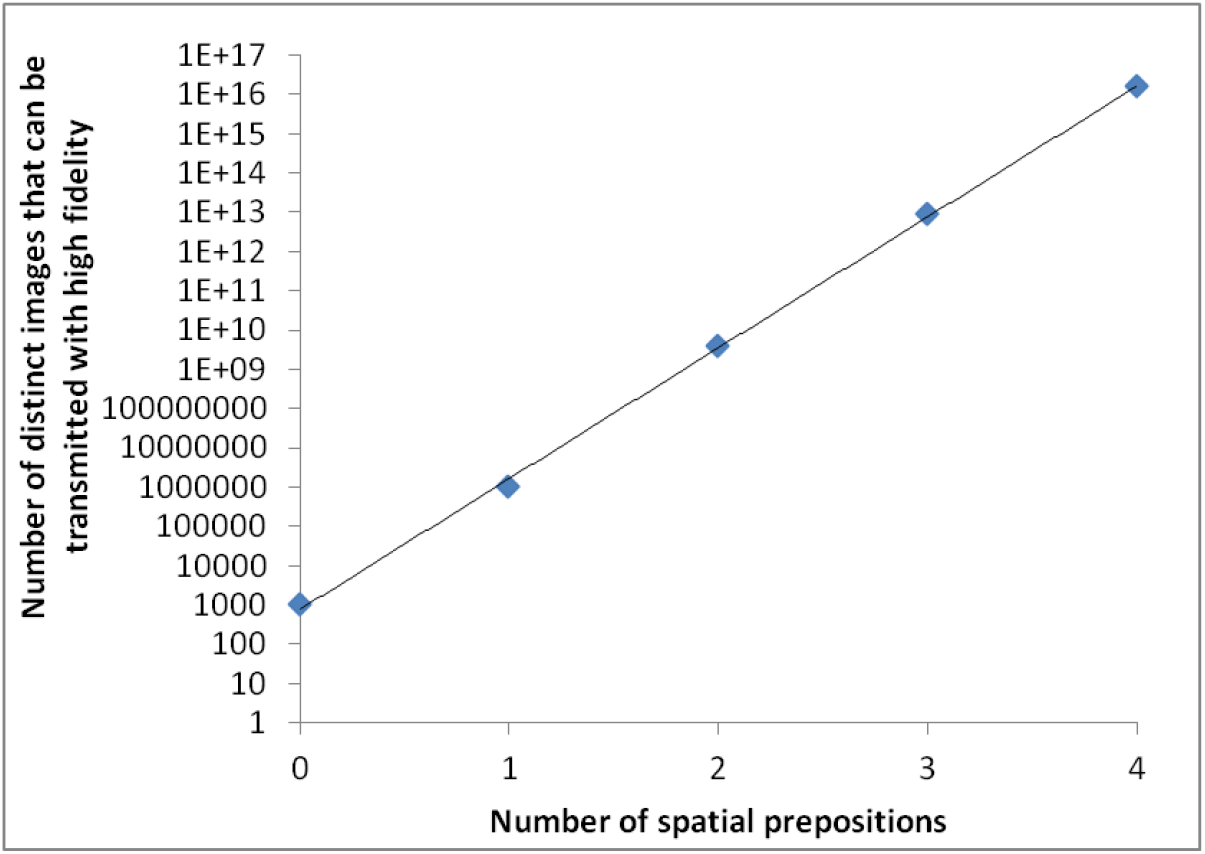
The graph shows the number of distinct objects combinations that can be conveyed with high fidelity in a communication system with 1,000 nouns as a function of the number of spatial prepositions.

The magic of using a finite number of words to communicate an infinite number of images completely depends on the PFS ability. A person lacking this ability is unable to construct novel mental images according to the rules imposed by spatial prepositions and, therefore, will not understand the meaning of spatial prepositions. In the theoretical example of a language with 1000 nouns and 100 spatial prepositions, the person with PFS disability will be limited in his/her comprehension to the 1000 nouns. Generalizing to other elements of language, we conclude that patients with PFS disability are expected to lack understanding of any recursive elements of language – the symptom reported by Fuster and Luria in *prefrontal aphasia* patients^15,18^. (Since “aphasia” is translated from Greek as “speechless” and these patients have no problem with speech, we prefer to call their condition a “PFS disability.”)

Extending this argument from a single individual to a community of individuals with PFS disability, we note that the communication system in that community must be non-recursive. Similarly, if we could envision a community of individuals who have not yet acquired the PFS ability phylogenetically, we could confidently say that they could not have understood recursive language and therefore could not have used spatial prepositions. They still could communicate, but their communication system must have been non-recursive, void of spatial prepositions and other sentences describing object combinations. Linguistics does not have an established name for such a communication system with thousands of words and no recursion. We cannot refer to it as ‘language’ in order to avoid confusion with recursive language. Accordingly, we will simply refer to it as rich-vocabulary non-recursive communication system.

### Use of recursive language in early childhood is necessary for acquisition of Prefrontal Synthesis

PFS disability is not limited to individuals with LPFC damage. Individuals without any brain injury exhibit PFS disability if they were not exposed to recursive language in early childhood^23–26^. In our metaanalysis of published research, ten out of ten individuals linguistically deprived until puberty suffered lifelong PFS disability despite learning significant vocabulary through intensive post-pubertal language therapy^4^. E.g., Genie, who was linguistically isolated until the age of 13 years 7 months, expanded her vocabulary to several hundred words following multi-year rehabilitation, but never completely acquired ability to understand spatial prepositions, verb tense, or recursion^27,28^. Furthermore, like other individuals with PFS disability, she invariably failed in nonverbal mental integration tasks that require PFS, such as those shown in Figure 1^4^. As a child Genie was not exposed to any dialogic communications; the next group of individuals grew up exposed to conversations, but not with a recursive language.

About 90% of all congenitally deaf children are born to hearing parents^29^. In the US, these children typically receive special services, are introduced to a formal sign language and are encouraged to use this recursive language for communication. In less developed countries, however, deaf children may never be exposed to a formal sign language. To communicate, such families usually spontaneously invent homesign (a.k.a. kitchensign), a system of iconic gestures that consists of simple signs. The homesign system though is lacking spatial prepositions, verb tenses and other recursive elements of a formal sign language. In other words, deaf linguistic isolates grow up exposed to a non-recursive communication system with large number of sign-words. Our analysis shows that these individuals deprived of recursive conversations until puberty performed poorly in all PFS tests, both verbal and nonverbal (such as those shown in Figure 1) despite focused multi-year post-pubertal rehabilitation efforts^4^. The consistent observation of PFS disability in these individuals stands in stark contrast to their performance on memory as well as semantic tests: they could easily remember hundreds of newly learned words and recall previously seen objects from memory but had real difficulty in any tasks requiring them to mentally combine these objects into novel configurations. Consider E.M., a deaf individual who was not introduced to a formal sign language until the age of 15^30^. E.M. tested at age 19, four years after his acquisition of hearing aids and intensive language therapy, could not follow a direction to “put the green box in the blue box”^30^; he would pick up the appropriate two boxes and “move them through a variety of spatial arrangements, watching the examiner for clues as to which was correct.”^30^ For successful completion of this task without relying on trial and error, one needs to use PFS to generate the novel mental image of the green box inside the blue box. In other words, the correct arrangement of physical objects is possible only after completing its mental simulation.

Isolated deaf children who grow up using homesign for communication must be distinguished from deaf children developing in a community of other deaf children, as they are known to be able to independently invent a recursive sign language of their own. In 1980, following the Sandinista revolution, the Nicaraguan government opened several vocational schools for deaf children. By 1983 there were over 400 students in the two schools. The school program emphasized spoken Spanish and lip reading, and discouraged the use of signs by teachers. The program failed and students were unable to learn the Spanish language in such a manner. However, the school provided fertile ground for deaf students to communicate with each other. In this process, children gradually spontaneously generated a new sign language, complete with *syntax,* verb agreement and other conventions of grammar^31–34^. Studying generational differences between Nicaraguan children who grew up when the sign language was in its initial stage of development and those who grew up a decade later exposed to a richer vocabulary and more complex recursive elements demonstrated clear cognitive differences between the different cohorts of children^23,24,26^.

Prelingual deafness is such a serious concern that the US government has enacted laws to identify affected newborns. In 1999, the US congress passed the “Newborn and Infant Hearing Screening and Intervention Act,” which gives grants to help states create hearing screening programs for newborns. Otoacoustic Emissions Testing is usually done at birth, followed by an Auditory Brain Stem Response if the Otoacoustic Emissions test results indicated possible hearing loss. Such screening allows parents to expose deaf children to a formal sign language as early as possible and therefore avoid any delay in introduction to recursive language.

Lack of communication with the use of recursive language is a big concern in children with autism^20^ and can lead to PFS disability. The autism community refers to the phenomenon whereby children cannot combine disparate objects into a novel mental image as *stimulus overselectivity,* or *tunnel vision,* or *lack of multi-cue responsivity*^35–37^. The ASD community is very aware of this problem and there is wide consensus that intense *early intervention* should be administered to children as soon as they are diagnosed with ASD^38^. The goals of speech language pathologists (SLP) and Applied Behavior Analysis (ABA) therapists happen to be built around acquisition of PFS. SLPs commonly refer to PFS developing techniques as “combining adjectives, location/orientation, color, and size with nouns,” “following directions with increasing complexity,” and “building the multiple features/clauses in the sentence”^39^. In ABA jargon, these techniques are known as “visual-visual and auditory-visual conditional discrimination”^40–43^, “development of multi-cue responsivity”^36^, and “reduction of stimulus overselectivity”^37^.

Despite the best efforts of therapists, 30-40% of individuals diagnosed with autism spectrum disorder (ASD) experience lifelong PFS disability and the associated impairment in the ability to understand spatial prepositions, verb tenses and recursion^44^. These individuals, commonly referred to as having low-functioning ASD, typically exhibit full-scale IQ below 70^20,45^ and usually perform below the score of 85 in non-verbal IQ tests^20^. In fact, the PFS ability and the associated understanding of spatial prepositions and recursion may be the most salient differentiator between high-functioning and low-functioning ASD.

The PFS ability develops in neurotypical individuals between the ages of three and five^19,46^, but the dependent relationship between the recursive dialogs and acquisition of PFS may already exist before the age of two. The randomized controlled study of institutionalized Romanian children demonstrated a significant difference at the age of eight between children placed in foster care and therefore exposed to recursive dialogs before the age of two and children who have been placed in foster care after the age of two. The former group performed better in mental integration tasks^47^ and showed increased myelination, hypothesized to be an important part of synchronous frontoposterior neural network essential for PFS^10^.

The next significant decline in plasticity is around the age of five. When the left hemisphere is surgically removed before the age of five (to treat cancer or epilepsy), patients often attain normal cognitive functions in adulthood (using the one remaining hemisphere). Conversely, removal of the left hemisphere after the age of five often results in significant impairment of recursive language and PFS-based cognitive skills^48–52^.

Further reduction of plasticity occurs by the time of puberty; a lack of experience in recursive dialogs before puberty invariably results in PFS disability^4^. While parts of the LPFC network retain some plasticity for a significantly longer period of time, since myelination of the LPFC continues into the third decade of life or later^53^, this plasticity seems inadequate to assist individuals who were deprived of recursive language until puberty in the acquisition of PFS despite many years of intensive language therapy^4^.

It is commonly accepted that childhood use of recursive language is essential for normal cognitive development^27,50,54,55^ and critical periods have been identified for many language-related functions, such as phoneme tuning^56,57^, grammar processing^58^, articulation control^59^, and vocabulary acquisition^60^. However, to understand language evolution, it is fundamentally important to realize the difference in critical periods for PFS and the rest of language-related functions. Phoneme tuning, grammar processing, articulation control, and vocabulary acquisition can all be significantly improved by training at any age^61,62^ and, therefore, have weak critical periods. PFS, on the other hand, has a strong critical period. Similar to other traits with strong critical periods, such as monocular deprivation^63^, filial imprinting in birds^64^, and monaural occlusion^65^, PFS cannot be acquired in adulthood. The neural infrastructure mediating PFS can only be established in early childhood.

### Evolutionary conundrum

As is the case with ontogenetic development of many neurological systems from muscle innervation to the development of all sensory systems, nature’s intent must be complemented by adequate nurture: normal development of vision requires stimulation of retina, normal development of hearing depends on auditory stimulation, normal development of somatosensory cortex is the function of tactile input, etc. What is highly unusual about the ontogenetic acquisition of PFS is that the necessary experience is provided by the exposure to a **purely cultural phenomenon**: dialogic communication using a recursive language.

For the normal development of vision, light reflected from surrounding objects has to reach the retina, but that occurs whenever it is light, independent of cultural exposure; for the normal development of the muscular system, the trophic factors released by muscles have to reach their neurons, but that occurs whenever a child is moving – the stimulation to neurons comes naturally even when a child is growing alone in a forest^66^. However, this is not the case with PFS. The development of neurological networks necessary for PFS in a child requires a community of humans using existing recursive language and willing to utilize it in communication with a child. Modern children who experience fewer conversational turns show significant reduction of frontoposterior fiber tracks^67^ and complete lack of recursive conversations is associated with complete PFS disability^4^.

Two observations have a profound consequence on phylogenetic acquisition of recursive language: (1) dialogs with non-recursive homesign systems do not suffice for the acquisition of PFS and (2) dialogs with the use to a recursive language have to occur during the period of highest neural plasticity, which peaks before the age of two, diminishes greatly after the age of five, and expires completely some time before puberty. Simply put, it is not enough to be fully genetically modern individual to acquire PFS, one needs to be exposed to recursive language early in childhood. This results in the proverbial ‘chicken and the egg’ problem since neither PFS nor recursive language could be acquired phylogenetically one before the other. This dependency creates an evolutionary barrier, which can be cleared only if multiple factors fall in place within a single generation. The following chapters put forward a hypothesis that resolves this conundrum by proposing that these two processes – the neurologically-based PFS and the culturally-transmitted recursive language – were acquired phylogenetically at the same time.

## The evolutionary context

### Acquisition of Prefrontal Synthesis around 70,000 years ago

Once we realize that PFS is a neurologically separate component of imagination, we must then ask when PFS was phylogenetically acquired^2^? Archeological records indicate gradual, piece-meal process of accretion of symbolic artifacts such as perforated shells^68^, use of pigments presumably in body decoration^68^, and intentional burials^69^, over hundreds of thousands of years^70^. However, symbolic thinking is not congruent to PFS. The symbolic use of objects can be accompanied by PFS in modern individuals, but PFS is not necessary for using an object as a symbol. For example, the use of red ochre may be highly symbolic due to its association with blood. However, this association may be entirely based on memory of an emotional event such as a bloody battle, as well as spontaneously formed imagery of a battle. Crucially, memory recall and spontaneously formed imagery (like in a dream or during an insight) do not rely on PFS^2^ and therefore use of red ochre is not an indication of the PFS abilities in hominins. Similarly, simple personal ornaments such as perforated shells^68,71–73^ could have been used as symbols of social power. However, neither their manufacturing nor their use signify the PFS ability. The line marks on stones and shells^74^, as well as geometrical figures and hand stencils painted on cave walls are undoubtedly associated with general improvement in the LPFC function and active imagination in their creators, but there is nothing in these artifacts indicating the presence of the most advanced component of active imagination, the PFS ability^2^.

The first definitive evidence of PFS appears in the archeological record around 65,000 to 40,000 and it emerges simultaneously in several modalities: (1) composite figurative arts, (2) bone needles with an eye, (3) construction of dwellings, and (4) elaborate burials. Together with (5) lightning-fast colonization of the globe and migration to Australia (presumably by boats) at around 62,000 years ago and (6) demise of the Pleistocene megafauna (presumably with the aid of animal traps) the archeological evidence indicates the presence of the PFS ability in hominins at about 62,000 years ago.

1. **Composite figurative objects**. Depiction of composite objects that don’t exist in nature provides an undeniable evidence of PFS. These composite objects such as the Lowenmensch (“lion-man”) sculpture from the caves of Lone valley in Germany (dated to 37,000 years ago)^75^ must have been imagined by the artists by first *mentally synthesizing* parts of the man and beast together and then executing the product of this mental creation in ivory or other materials. The composite artworks such as lion-man from Germany, a bird-man from Lascaux, a lion-woman from Chauvet, and the engraving of a bird-horse-man from Hornos de la Peña provide a direct evidence that by 37,000 years ago some humans were capable of PFS.
2. **Creativity and innovation**. Improvement of stone tools is our best indication of improving active imagination. Turning an unformed cobblestone into a sharp tool requires an active purposeful imagination of a previously unseen object. According to Ian Tattersall, “To make a carefully shaped handaxe from a lump of rock not only demanded a sophisticated appreciation of how stone can be fashioned by fracture, but a mental template in the mind of the toolmaker that determined the eventual form of the tool”^76^. This “mental template” is different from the original cobblestone and therefore could not have been recalled from memory. It must have been actively imagined by the toolmaker. Apes do not manufacture stone tools in the wild and attempts to teach stone tools manufacturing to apes have failed^77^, suggesting that this ability was acquired after humans have split from chimpanzee line 6 million years ago. The first stone tools, Mode One stone choppers, dated to about 3.3^78^ to 2.5^79^ million years ago are crude and asymmetrical. Starting from about 2 million years ago, hominins were capable of manufacturing a fine symmetrical Mode Two handaxes^69^. Neanderthals manufactured even better Mode Three Mousterian tools found in the archeological record from about 0.4 million years ago^69^. If the quality of stone tools is informing us of the quality of mental template and the corresponding LPFC ability to mold their percept into the mental template, then stone tools provide a time record of active imagination improving over the last 3 million years. However, general improvement in active imagination is not informing us on the presence of PFS. Modern individuals with PFS disability can manufacture any of the stone tools (unpublished observations) or wooden spears, suggesting that less sophisticated components of imagination can suffice for stone tools manufacturing process. In fact, individuals with severe intellectual disability (majority of whom have PFS disability) can be organized to manufacture all kinds of crafts^80^. What tools can unambiguously signify the presence of PFS in hominins? One of the most obvious examples of tools clearly associated with PFS are bone needles used for stitching clothing. To cut and stitch an animal hide into a well-fitting garment, one needs first to mentally simulate the process, i.e. imagine how the parts can be combined into a finished product that fits the body. Such mental simulation is impossible without PFS. Earliest bone needles are dated to 61,000 years ago^81^ and they provide the first unambiguous indication of PFS in some individuals.
3. **Design and construction**. Human dwellings are not built by reflex. An integral part of design and construction is visual planning, which relies on the mental simulation of all the necessary construction steps, which is impossible without PFS. There is little evidence of hominins constructing dwellings or fire hearths until the arrival of *Homo sapiens.* While Neanderthals controlled the use of fire, their hearths were usually very simple: most were just shallow depressions in the ground. There is almost a complete lack of evidence of any dwelling construction at this period^82^. Conversely, the arrival of *Homo sapiens* is marked by a multitude of constructed structures including stone-lined and dug-out fireplaces, as well as unambiguous remains of dwellings, which all flourished starting around 30,000 years ago. These include foundations for circular hut structures at Vigne-Brune (Villerest) in eastern France, dating back to 27,000 years ago^83^; postholes and pit clusters at a site near the village of Dolní Věstonice in the Czech Republic, dating back to 26,000 years ago^84^, and mammoth bone structures at Kostienki, Russia and Mezirich, Ukraine^85^.
4. **Adorned burials and religious beliefs**. Religious beliefs and beliefs in afterlife are the ultimate products of PFS. An individual with PFS disability cannot be induced into believing in spirits, as they cannot imagine gods, cyclops, mermaids, or any other mythological creatures. The origin of religious beliefs can be traced by following the evidence of beliefs in the afterlife. Beliefs in the afterlife, in turn, are thought to be associated with adorned burials. Therefore, the development of religious beliefs may be inferred by studying the time period when humans started to bury their deceased in elaborate graves with accompanying “grave goods.” The oldest known human burial, dated at 500,000 years ago and attributed to *Homo heidelbergensis,* was found in the Sima de los Huesos site in Atapuerca, Spain, and consists of various corpses deposited in a vertical shaft^86^. A significant number of burials are also associated with Neanderthals: La Chapelle-aux-Saints, La Ferrassie, and Saint-Cesaire in France; Teshik-Tash in Uzbekistan; Shanidar Cave in Iraq^87^. However, whether or not these sites constitute actual burial sites is hotly disputed. Their preservation could well be explained by natural depositions^88^. Even if those burials were made deliberately, the goal may have been to simply toss the bodies away in order to discourage hyena intrusion into the caves^76^. In any case, these early burials completely lack the “grave goods” that would indicate the belief in an afterlife^76^. Human skeletal remains that were intentionally stained with red ochre were discovered in the Skhul and Qafzeh caves, in Levant and dated to approximately 100,000 years ago^89^. One of the burials contains a skeleton with a mandible of a wild boar, another contains a woman with a small child at her feet, and yet another one contains a young man with a possible offering of deer antlers and red ochre^90^. While these burials are clearly intentional, whether or not they indicate the belief in an afterlife is uncertain. The ochre by itself is an inconclusive evidence. For example, ochre could have been used during lifetime to protect skin from insects^91^ and the deceased could have been buried still bearing the ochre marks. The small number of “offerings” found in these burial sites may have simply been objects that fell into the burial pit accidentally. In any case, there is not enough conclusive evidence from these early burials to judge the occupants’ beliefs in an afterlife. The number of known *adorned* burials and the sophistication of the offerings significantly increase around 40,000 years ago. To date, over one hundred graves of *Homo sapiens* have been discovered that date back to the period between 42,000 and 20,000 years ago^92^. In many cases several bodies were interred in a single grave. Burial offerings were commonplace and ochre was used abundantly. Examples include: a burial in Lake Mungo, Australia of a man sprinkled with red ochre, dating back to 42,000 years ago^93^; an elaborate burial in Sungir, Russia that includes two juveniles and an adult male wearing a tunic adorned with beads and carefully interred with an astonishing variety of decorative and useful objects, dating back to 30,000 years ago^94^; a grave in Grimaldi, Italy, which contains the remains of a man and two adolescents along with burial offerings from around 40,000 years ago^92^; and a site in Dolni Vestonice, in the Czech Republic where a woman was buried between two men and all three skulls were covered in ochre dating back to 28,000 years ago^95^. The appearance of adorned burials in multiple geographical locations is consistent with the PFS ability in most individuals by 40,000 years ago^92^.
5. **Fast colonization of the globe and migration to Australia**. Hominins diffusing out of Africa had been colonizing the Europe and Asia long before the arrival of *Homo sapiens:* the remains of *Homo erectus* have been found as far as in Spain^96^ and Indonesia^97^ and Neanderthals remains have been found in Europe and Asia^98^. However, both the extent and the speed of colonization of the planet by *Homo sapiens* 70,000 to 65,000 years ago are unprecedented. Our ancestors quickly settled Europe and Asia and crossed open water to Andaman Islands in the Indian Ocean by 65,000 years ago^99^ and Australia as early as 62,000 years ago^100^. Migration to Australia is consistent with the use of boats by early modern humans further underlying their propensity for technological innovations.
6. **Building animal traps and demise of the Pleistocene megafauna**. Without PFS one cannot envision the building of an animal trap, e.g. pitfall trap, which requires digging a deep pit and camouflaging it with twigs and branches. PFS aids trap building in three ways. First, a leader can use PFS to mentally simulate multiple ways to build a trap. Second, a leader could use PFS to think through the step-by-step process of building a trap. Finally, a leader could communicate the plan to the tribe: “We will make a trap by digging a large pit and covering it with tree branches. A mammoth will then fall into the pit; no need to attack a mammoth head on.” In fact, early modern humans are known for building traps; traps for herding gazelle, ibex, wild asses and other large animals were found in the deserts of the Near East. Some of the traps were as large as 60km (37miles) in length^101^. Funnel-shaped traps comprising two long stone walls (up to 60 kilometers in length!) converged on an enclosure or pit at the apex. Animals were probably herded into the funnel until they reached the enclosure at the apex surrounded by pits, at which point the animals were trapped and killed. Some traps date back to as early as the 7th millennium BC^101^. The building process must have been pre-planned by a tribe leader (or several leaders) and then explained to all the workers. Each worker, in turn, would have had to understand exactly what they needed to do: collect proper stones, assemble stones into a wall, and have the two walls meet at the apex 60 km away from where they started. The correlation of human migration with demise of the Pleistocene megafauna^102,103^ is consistent with PFS that would have enabled mental planning of sophisticated attack strategies with the use of animal traps^101^.

#### Conclusions from paleontological evidence

There is no evidence of the PFS ability in hominins before 65,000 years ago and there is an *abundance* of clear and unambiguous evidence of the PFS ability in hominins after around 62,000 years ago. Composite objects executed in bone and cave paintings, bone needles with an eye, construction of dwellings, appearance of adorned burials, and steadfast colonization of the planet are all the external manifestations of PFS. The PFS-related artifacts are highly correlated with each other in time and geography and are associated with *Homo sapiens* diffusion out of Africa around 70,000 years ago. This abrupt change toward modern imagination has been characterized by paleoanthropologists as the “Upper Paleolithic Revolution,”^104^ the “Cognitive revolution,”^105^ and the “Great Leap Forward”^106^ and it is consistent with acquisition of PFS sometime shortly before 62,000 years ago (for a more skeptical position, see^70^. Remember, however, that researchers arguing for a more gradual cultural and technological elaboration do not differentiate between ‘symbolic artifacts,’ general ‘active imagination artifacts,’ and specific ‘PFS artifacts.’ There is no doubt that accretion of ‘symbolic artifacts’ (use of ochre) and ‘active imagination artifacts’ (stone tools) is gradual over hundreds of millennia. It is the appearance of specific PFS evidence that seems to be relatively abrupt). The genetic bottleneck that has been detected around 70,000^107^ may have been associated with “founder effect” of few individuals who acquired PFS and thus developed a significant competitive advantage over the rest of hominins. In the rest of the manuscript we try to deduce what could have triggered their PFS acquisition.

### Could development of articulate speech trigger prefrontal synthesis acquisition 70,000 years ago?

The articulate speech of humans is unique among primates. The vocal tract of our closest relatives, chimpanzees, is extremely limited in its ability to modulate sound. While there is no theoretical limit on the number of different vocalizations nonhuman primates can generate^108^, attempts to teach chimpanzees articulate speech have failed^109^ and the range of distinct vocalizations observed in the wild is limited to between 20 and 100^110–113^. On the contrary, human languages contain tens of thousands of different words easily generated by the modern human vocal apparatus. If development of articulate speech could have triggered PFS acquisition, that would explain the human cognitive revolution 70,000 years ago. Unfortunately, the dates do not match.

Evolutionary changes in the vocal tract have been extensively studied by paleoanthropologists^76,114,115^. The modern vocal apparatus developed as a result of changes of the structure and the position of many organs that play a role in generating and modulating vocalizations: larynx, tongue, musculature of the mouth, lips, and diaphragm as well as the neurological control of the associated musculature.

While cartilaginous and soft tissue is not preserved in the fossil record, we can draw conclusions about the evolution of vocal apparatus from the bony structures which do survive. Dediu and Levinson cite five lines of converging evidence pointing to acquisition of modern speech apparatus by 600,000 years ago^1^: (1) the changes in hyoid bone, (2) the flexion of the bones of the skull base, (3) increased voluntary control of the muscles of the diaphragm, (4) anatomy of external and middle ear, and (5) the evolution of the FOXP2 gene.

1. **The changes in hyoid bone**. This small U-shaped bone lies in the front of the neck between the chin and the thyroid cartilage. The hyoid does not contact any other bone. Rather, it is connected by tendons to the musculature of the tongue, and the lower jaw above, the larynx below, and the epiglottis and pharynx behind. The hyoid aids in tongue movement used for swallowing and sound production. Accordingly, phylogenetic changes in the shape of the hyoid provide information on the evolution of the vocal apparatus. The hyoid bone of a chimpanzee is very different from that of a modern human^116^. The australopith hyoid bone discovered in Dikika, Ethiopia, and dated to 3.3 million years ago closely resembles that of a chimpanzee^117^. The *Homo erectus* hyoid bone recovered at Castel di Guido, Italy, and dated to about 400,000 years ago reveals the “bar-shaped morphology characteristic of *Homo,* in contrast to the bulla-shaped body morphology of African apes and *Australopithecus”*^118^. Neanderthal hyoids are essentially identical to that of a modern human in size and shape: these have been identified in Kebara, Israel and^119^ El Sidrón, Spain^120^. At the same time these are also identical to hyoid of *Homo heidelbergensis* from Sima de los Huesos, Spain^121^ suggesting that the latter was a direct ancestor of both *Homo neanderthalensis* and *Homo sapiens* and had already possessed a nearly modern hyoid bone^1,122^. The similarities between Neanderthal and modern human hyoid make it likely that the position and connections of the hyoid and larynx were also similar between the two groups.
2. **The flexion of the bones of the skull base**. Laitman^123,124^ has observed that the roof of the vocal tract is also the base of the skull and suggested that evolving vocal tract is reflected in the degree of curvature of the underside of the base of the skull (called basicranial flexion). The skull of *Australopithecus africanus* dated to 3 million years ago shows no flexing of the basicranium, as is the case with chimpanzees^125^. The first evidence of increased curvature of the base of the basicranium is displayed in *Homo erectus* from Koobi Fora, Kenya, 1.75 million years ago^123^. A fully flexed, modern-like, basicranium is found in several specimen of *Homo heidelbergensis* from Ethiopia, Broken Hill 1, and Petralona from about 600,000 years ago^126^.
3. **Increased voluntary control of respiratory muscles**. Voluntary cortical control of respiratory muscles is a crucial prerequisite for complex speech production^127^. Greater cortical control is associated with additional enervation of the diaphragm, that can be detected in fossils as an enlarged thoracic vertebral canal. *Homo erectus* from 1.5 million years ago (Turkana Boy) has no such enlarged canal, but both modern humans and Neanderthals do^1^, providing converging evidence for acquisition of modern-like vocal apparatus by 600,000 years ago.
4. **The anatomy of external and middle ear**. Modern humans show increased sensitivity to sounds between 1kHz and 6kHz and particularly between 2kHz and 4kHz. Chimpanzees, on the hand, are not particularly sensitive to sounds in this range^121,128^. Since species using complex auditory communication systems, tend to match their broadcast frequencies and the tuning of perceptual acuity^129^, it was argued that changes in the anatomy of external and middle ear in hominins are indicative of the developing speech apparatus. Data from several Neanderthal and *Homo heidelbergensis* fossils indicate a modern-human like pattern of sound perception with highest sensitivity in the region around 4kHz, that is significantly different from that of chimpanzees^128,130^.
5. **The evolution of the FOXP2 gene**. The most convincing evidence for the timing of the acquisition of the modern speech apparatus is provided by DNA analysis. The FOXP2 gene is the first identified gene that, when mutated, causes a specific language deficit in humans. Patients with FOXP2 mutations exhibit great difficulties in controlling their facial movements, as well as with reading, writing, grammar, and oral comprehension^131^.The protein encoded by the FOXP2 gene is a transcription factor. It regulates genes involved in the production of many different proteins. The FOXP2 protein sequence is highly conserved. There is only one amino acid difference in the chimpanzee lineage going back some 70 million years to the common ancestor with the mouse^132^. The FOXP2 proteins of chimpanzee, gorilla and rhesus macaque are all identical. This resistance to change suggests that FOXP2 is extraordinarily important for vertebrate development and survival. Interestingly, there is a change of two amino acids in FOXP2 that occurred over the last 6 million years, during the time when the human lineage had split off from the chimpanzee. These two amino acid substitutions predate the human-Neanderthal split. Both amino acid substitutions were found in two Neanderthals from Spain^133^, as well as in Neanderthals from Croatia^134^, and in Denisovans, an extinct Asian hominin group related to Neanderthals^135^. This indicates that *Homo heidelbergensis,* the common ancestor of *Homo sapiens* and Neanderthals, already had the two “human specific” amino acid substitutions. Despite evidence of possible further evolution of FOXP2 in *Homo sapiens*^136^, the comparatively fast mutation rate of FOXP2 in hominins indicates that there was strong evolutionary pressure on development of the speech apparatus before *Homo sapiens* diverged from Neanderthals over 500,000 years ago^137^.

#### Conclusions on acquisition of articulate speech

Based on these five lines of evidence — the structure of the hyoid bone, the flexion of the bones of the skull base, increased voluntary control of the muscles of the diaphragm, anatomy of external and middle ear, and the FOXP2 gene evolution — most paleoanthropologists conclude that the speech apparatus experienced significant development starting with *Homo erectus* about two million years ago and that it reached modern or nearly modern configurations in *Homo heidelbergensis* about 600,000 year ago^176^. Dediu and Levinson write: “there is ample evidence of systematic adaptation of the vocal apparatus to speech, and we have shown that this was more or less in place by half a million years ago”^1^. We will never know the extent of *Homo heidelbergensis* neurological control of their speech, however, considering that chimpanzee communication system already has 20 to 100 different vocalizations^110–113^, it is likely that the modern-like remodeling of the vocal apparatus in *Homo heidelbergensis* extended their range of vocalizations by orders of magnitude. In other words, by 600,000 years ago the number of distinct verbalizations used by hominins for communication was on par with the number of words in modern languages. Thus, by 600,0 years ago the number of words in the lexicon could NOT have been holding back acquisition of PFS and recursive language. Articulate speech likely has been an important prerequisite to, but could not be the trigger for PFS acquisition 70,000 years ago. It also follows that hominin groups with fluent articulate speech must have existed for hundreds of millennia before acquisition of PFS. In many regards these hominins must have been similar to patients with prefrontal aphasia discussed in the introduction who have fluent speech, but completely limited to non-recursive dialogs due to PFS disability.

### Young children must have invented first recursive elements of language – the Romulus and Remus hypothesis

As discussed in the introduction, PFS is not acquired ontogenetically unless children are exposed to recursive language in early childhood. According to our analysis, non-recursive communication system (called kitchensign or homesign, as opposed to a formal sign language) is unable to facilitate acquisition of PFS even in genetically modern children^4^. These children uniformly exhibit lifelong PFS disability as assessed by both verbal and nonverbal tests despite many years of focused post-pubertal rehabilitation. Our analysis shows that early childhood use of recursive language is essential for acquisition of PFS^4^. Thus, it follows that phylogenetically, PFS must have been acquired at the same time as recursive language. Since PFS was likely acquired around 70,000 years ago, we can only assume that recursive language was also acquired at the same time.

Furthermore, since only children can acquire PFS, it follows that around 70,000 years ago young children must have invented the first recursive language. The parents of these children used a rich-vocabulary communication system for millennia. That system, however, contained no spatial prepositions, nesting, verb tenses or other recursive elements of language. The children may have stumbled upon recursive elements of language such as spatial prepositions (development of new words and even complete language is a common phenomena among very young children living together, the process called cryptophasia^138^). With just a few spatial prepositions, their communication system would be able to communicate nearly infinite number of novel images (Figure 2) and therefore their dialogs would have provided enough stimulation to acquire PFS^4^. Accordingly, we named our hypothesis after the celebrated twin founders of Rome, Romulus and Remus. Similar to legendary Romulus and Remus whose caregiver was a wolf, the real children’s caregivers had an animal-like communication system with many words but no recursion. These children were in a situation reminiscent of the condition of the children who invented the Nicaraguan Sign Language: their parents could not have taught them spatial prepositions or recursion; children had to invent recursive elements of language themselves. We can expect that each following generation expanded the recursive elements of language and, as a result, improved their PFS. Such parallel development of newly invented language and PFS is found among deaf children in Nicaragua. As newer generations of Nicaraguan Sign Language speakers expanded their language, they have also improved on multiple measures related to PFS^23,24,26^.

### The genetic trigger

The Romulus and Remus hypothesis attempts to explain the more than 500,000-year gap between acquisition of modern speech apparatus and recursive language by a low probability of an event when two or more very young children living together concurrently (1) invent recursive elements of language, (2) have enough dialogs to stimulate each other’s acquisition of PFS, and (3) survive to adulthood to take advantage of their modern behavior and procreate. Unfortunately, in its pure form, the Romulus and Remus hypothesis does not survive a simple numerical test. A hominin tribe of 150 individuals spaced linearly from 0 to 30, has 5 peers. Even if we assume (1) that children younger than two could not invent any new words since they did not speak articulately enough, (2) children needed at least a year of using recursive dialogs to stimulate the formation of the neurological networks responsible for PFS, and (3) the end of the critical period for PFS acquisition occurs at the age of five, the model still yields a group of 15 children from two to five years of age per tribe. Fifteen children at the peak of their plasticity is on par with the number of deaf students, who spontaneously invented the Nicaraguan sign language (400 students in two schools)^31–34^. It is hard to explain why 15 children in any of the many hominin tribes have not invented recursive language in over 500,000-year period, given that they already had nonrecursive communication system and only had to invent recursive elements, while the Nicaraguan deaf children invented both in a few generations.

To further refine understanding of the number of children, an evolutionary mathematical model of a hominin tribe was generated based on the Australian aboriginal population. Moody^139^ reported that Australian aboriginal children experienced disproportionally higher mortality than adults with at least 28% of second-year deaths, and about 9% of deaths in the two to four years age group. This pattern of childhood mortality is best described by an exponential function of age (Mortality=const+0.4*EXP(-age/2)). After the const was calibrated to generate a stable tribe population, the model predicted the total of 207 tribesmen (100 individuals younger than 12 and 107 individuals 12 and older). Importantly the model predicted 25 children age 2 to 5, i.e. satisfying the conditions (1) to (3). Thus, the population model demonstrates that it is impossible to explain the 500,000-year gap between acquisition of modern speech apparatus and PFS by a cultural process of invention of recursive elements of language alone. Even under strict conditions (1) to (3) **a genetically modern tribe is expected to invent recursive elements and acquire PFS within several generations similar to the Nicaraguan deaf community**. It follows that PFS acquisition must have been also precluded by a genetic factor.

What may have been the genetic difference that prevented children from inventing recursive elements of language and acquiring PFS for hundreds of millennia? Inadequate vocal apparatus is commonly brought up to explain the conundrum. However, as discussed above, the improvements to the vocal apparatus amassed by hominins by 600,000 years ago must have increased vocabulary by several orders of magnitude, from 20 to 100 in chimpanzees^110–113^ to thousands of different words in hominins. Modern children start acquiring PFS at the age of three^19^, while using no more than few hundred words^140^ and, therefore, **the number of words spoken by hominins 600,000 years ago could not have been the limiting factor to acquisition of PFS**.

From a neuroscience perspective, it is relatively easy to imagine how a single mutation could have increased the brain volume, or the number of synapses, or the number of glial cells, or the extent of axonal myelination by a few percentage points, but those relatively small neurological differences could not have prevented children from PFS acquisition. The one neurological difference that could have a direct effect on PFS acquisition is the duration of critical period. If the duration of critical period in pre-PFS hominins was shorter than in modern children, that would have decreased the probability of invention of recursive elements and at the same time having enough time to train their dialog-dependent neurological networks essential for PFS^67^. For example, if the critical period for acquisition of PFS was over by the age of two, hominin children would have no chances for acquiring PFS at all. Only a critical period ending at the age of three would have provided a minimal opportunity to acquire PFS.

The duration of critical period for PFS acquisition is unknown in hominins, but has been tested in apes, first in terms of language acquisition and second in terms of the rate of PFC development. In many experiments, apes were raised in human environment and exposed to recursive language from infancy. These animals commonly learn hundreds to thousands of words but never acquired PFS (tested linguistically and nonverbally)^141–144^, consistent with some neurobiological barrier, possibly, a short critical period, preventing them from acquiring PFS. Additionally, genetic and imaging studies showed that ape’s PFC develops significantly faster than PFC in modern humans. The peak of synaptogenesis in the chimpanzee and macaque PFC occurs during several postnatal months, whereas in the human PFC it is shifted to about 5 years of age^145,146^. Similarly, the PFC myelination rate in chimpanzees is significantly faster than in humans^147^. Thus, in human children the PFC remains immature with respect to synaptogenesis for significantly longer period compared to chimpanzees and macaques.

Overall, humans are born with a less mature brain and develop 1.5-2 times slower than chimpanzees: molar teeth erupt three years later and humans become sexually active roughly five years after the chimps do^148^. However, the delay in maturation in the PFC from a few months in chimpanzees and macaques to five years in humans is much more dramatic compared to this overall delay in maturation. Additionally, this delay is exhibited by the PFC, but not by the cerebellum^149^. Sometime during the past six million years one or several genetic mutations have fixed in the human population causing this remarkable delay of the PFC maturation schedule. Notably, Liu *et al.* report that the delay in the PFC development occurred within the last 300,000 years, placing the “PFC delay” mutation on the radar for mutations that could have triggered the “Great leap forward” 70,000 years ago^145^. By slowing down PFC development this mutation could have prolonged critical period and enabled children’s invention of recursive elements, resulting in recursive dialogs and acquisition of PFS (similar to the Nicaraguan deaf children).

### It is likely that “PFC delay” and PFS were acquired simultaneously

Mutations that get selected and fixed in a population are usually associated with some survival benefits (e.g., lactase persistence is associated with the continued ability to digest lactose in milk after weaning). Mutations that decrease an organism’s ability to survive and reproduce are selected against and not passed on to future generations. The “PFC delay” mutation is inherently a strange mutation. By prolonging the critical period, the “PFC delay” mutation increases the chances for acquisition of PFS. At the same time, this mutation carries clear disadvantages. A decrease in the PFC development rate results in a prolonged immaturity when the brain is incapable of full risk assessment. For example, three-year-old chimps often venture away from their mother, but rarely come close to water, their decision-making PFC prohibiting them from doing so. On the contrary, in human children under 4 years of age, drowning is the leading cause of mortality, resulting in over 140,000 deaths a year^150^. The PFC of the four-year-old child is unable to fully assess the risk of drowning. Similarly, three-year-old children cannot be left alone near fire, near an open apartment window, near a traffic road, or in a forest. **In terms of risk assessment, three-year-old humans are intellectually disabled compared to any other three-year-old animal**. From the point of view of risk assessment, an individual with slower PFC maturation rate has lower chances to survive childhood, unless risks are mitigated by culture (e.g., we hold small children by hand near roads and cliffs, buckle them in a high chair, and never let them outside alone). Culture, however, could not have immediately caught up to delayed PFC maturation. Thus, at least initially “PFC delay” is expected to increase childhood mortality.

The evolutionary mathematical model was used to study the effect of increased childhood mortality due to PFC maturation slowdown. Decreasing childhood survival by 10% results in the collapse of tribe population to 50% within 150 years. If the “PFC delay” mutation did not lead to an immediate survival benefit that could have balanced the increase of childhood mortality, it would be expected to be weeded out from a hominin population.

How much of a post-pubertal benefit, the “PFC delay” mutation must have resulted in order to balance out the 10% increase in childhood mortality? The population model predicts that a minimum of 26% of post-pubertal increase in survival rate is required to keep a stable tribe population. Slowing down PFC maturation could have theoretically improved PFC-mediated social behavior, working memory, and impulse control, but it is hard to see how these traits could have increased adult survival by 26%. On the other hand, acquisition of PFS with its associated dramatic improvement in hunting enabled by animal trapping, stratagem, and new weaponry can easily explain this dramatic increase in adult survival. We conclude that it is highly unlikely that the “PFC delay” trait have evolved for some other function, persisted in a population, and later, after many generations, was adapted for PFS acquisition. In order to balance the immediate increase in childhood mortality associated with delayed PFC development, PFS acquisition must have quickly followed “PFC delay” acquisition.

Two or more young children with “PFC delay” must have been born at the same time and lived together for several years, so that they could often talk to each other (Box 1). These children were in a situation reminiscent of the condition of the youngsters who invented the Nicaraguan Sign Language: they were genetically modern (in terms of the “PFC delay” mutation), but their parents could not have taught them spatial prepositions; children had to invent recursive elements of language themselves. Having invented recursive elements and having acquired PFS, these children would have gained near-modern imagination: a ticket to dramatically improved hunting by trapping animals, proclivity for fast discovery of new tools through mental simulations and the ability to strategize over clever ways to eliminate other hominin competitors. The “PFC delay” mutation and recursive language could have then spread like a wildfire through other *Homo sapiens* tribes carried by new weapons and an elaborate stratagem made possible by the new recursive language. Improved survival as a result of burgeoning diet and comfortable hunting style can easily explain the unprecedented explosion of human population at the end of the upper Paleolithic^104^.

## Discussion

In this manuscript we presented a Romulus and Remus hypothesis of language acquisition that attempts to explain more than 500,000-year-long gap between the emergence of modern speech apparatus and the abundant evidence of modern imagination 70,000 years ago. We proposed that despite acquisition of modern vocal apparatus by 600,000 years ago, hominin communication system remained non-recursive. Spatial prepositions, verb tenses, and nesting were missing from their communication system not because there were not enough distinct vocalizations, but due to limitations of hominins’ imagination.

### Language is always limited by imagination

No animal can ever be taught to follow a command to bring a *‘long red* straw’ placed among several decoy objects including other *red* shapes (Lego pieces, small *red* animals) and *long/short* straws of other colors, not because animals cannot learn words for colors, sizes, or objects, but since they cannot purposefully imagine an object in different colors and sizes^2^.

This is not to say that animals cannot imagine objects in different colors or sizes spontaneously or in their dreams. Spontaneous imagination, however, is completely different neurobiologically from active purposeful imagination^2^. Active purposeful imagination is always driven by the LPFC and mediated by the frontoposterior fibers, while spontaneous imagination is independent of the LPFC.

Many scientists make a mistake of assuming human-like imagination in hominins. In the past, many people even extended human-like imagination to animals (e.g., St. Francis preaching to the birds). Anthropomorphism is the natural intuitive fallback for the unknown, but it has to be removed from science based on experimental evidence. Imagination is not a single phenomenon but includes multiple neurobiologically distinct mechanisms^2^. This insight, clear to most therapists working to build up active imagination mechanisms one by one in children with ASD, has been an enigma to many evolutionary biologists who measure hominin mind abilities through introspection.

Once we realize the existence of multiple neurobiologically distinct mechanisms of imagination, the natural question is when these mechanisms were phylogenetically acquired? Obviously, all the distinct mechanisms of imagination could not have been acquired at the same time. The best indication of improving active imagination in hominins is provided by the stone tools record as turning an unformed cobblestone into a sharp tool requires an active purposeful imagination of a previously unseen object^2^. The quality of stone tools that improved dramatically over the last 3 million years, from crude Mode One stone choppers, dated to 3.3 million years ago^78^, to symmetrical Mode Two handaxes manufactured from 2 million years ago^69^, to Mode Three tools manufactured from 0.4 million years ago^69^ provides the time record of the increasing ability of the LPFC to control the imaginary percept, but does not inform us on the Prefrontal Synthesis (PFS) ability. PFS requires significantly more complex neurobiological organization than that necessary for stone tools manufacturing and integration of modifiers^2^. PFS is the ultimate ability of modern humans, the measure stick of modern imagination, behavior, and culture. Unequivocal PFS evidence is completely missing from the archeological evidence before 70,000 years ago but is abundantly present after 62,000 years ago. Clear PFS evidence include (1) composite figurative arts, (2) bone needles with an eye, (3) construction of dwellings, and (4) elaborate burials. Together with (5) exceptionally fast colonization of the globe and migration to Australia (presumably by boats) at around 62,000 years ago and (6) demise of the Pleistocene megafauna (presumably with the aid of animal traps) this multitude of the archeological evidence indicates acquisition of PFS by some individuals around 70,000 years ago and their relentless conquest of the planet.

Since PFS is essential for comprehension of spatial prepositions and recursion, recursive modern-looking language could not have been acquired before PFS acquisition 70,000 years ago. We explained the 500,000 year-long period between acquisition of a modern speech apparatus and recursive language by existence of two evolutionary barriers associated with a critical period for PFS acquisition. One barrier is cultural, the other is genetic. Conversations with the use of recursive language provide an essential training for formation of frontoposterior connections necessary for PFS^67^. These connections only develop as a result of experience provided in early childhood by recursive language. Modern children who experience fewer conversational turns show significant reduction of frontoposterior fiber tracks^67^ and complete lack of recursive dialogs is associated with complete PFS disability^4^. Since hominin children were not involved in recursive conversations, they did not acquire PFS and, therefore, as adults, could not learn recursive language. Consequently, they could not teach recursive language to their own children, who, therefore, were not exposed to recursive conversations, continuing the cycle.

Even in genetically modern humans, this dependence of PFS on recursive language and recursive language on PFS creates an unsurpassable cultural barrier in isolated individuals. Modern children who are not exposed to recursive conversations during the critical period do not invent recursive language on their own and, as a result, never acquire PFS^4^. Considerable accumulation of young children in one place for several years is required to spontaneously invent recursive language (400 children in two schools in Nicaragua^31–34^).

The second evolutionary barrier to acquiring recursive language could have been a faster PFC maturation rate and, consequently, a shorter critical period. In modern children the critical period for PFS acquisition closes around the age of 5^4^. If the critical period in hominin children was over by the age of two, they would have no chance acquiring PFS. A longer critical period is imperative to provide enough time to both invent recursive language and train PFS via recursive conversations. We conjectured that the “PFC delay” mutation that is found in all modern humans, but not in Neanderthals^145,146^ was that last piece of necessary genetic makeup and that it triggered simultaneous synergistic acquisition of PFS and recursive language. The “Romulus and Remus” hypothesis calls for (1) two or more children with extended critical period due to “PFC delay” mutation; (2) these children spending a lot of time talking to each other; (3) inventing the recursive elements of language, such as spatial prepositions; (4) acquiring recursive-dialog-dependent PFS; and (5) surviving to adulthood and spreading their genes and recursive language to their offsprings. As adults, Romulus and Remus could immediately entertain the benefits of the newly acquired mental powers. They could have engineered better weapons and plan a sophisticated attack strategy using animal traps and stratagem. They would have become more successful builders and hunters and quickly reach the position of power enabling them to spread their genes more efficiently. The genetic bottleneck that has been detected around 70,000^107^ may have been associated with “founder effect” of a few individuals who acquired PFS and nearly completely replaced the rest of hominins.

### A comparison of the Romulus and Remus hypothesis to other theories

There are clear similarities of Romulus and Remus hypothesis to Chomsky’s Prometheus. Chomsky suggested that “roughly 100,000+ years ago, … a rewiring of the brain took place in some individual, call him Prometheus, yielding the operation of unbounded Merge, applying to concepts with intricate (and little understood) properties.”^151^. We argue however, that Prometheus could not have evolved alone. Modern children who are not exposed to recursive language before puberty cannot acquire PFS later in life^4^. If parents did not expose Prometheus to recursive language, the only way for Prometheus to acquire PFS was to invent recursive language himself and then use it to train his own dialog-dependent PFS. This fit can only be accomplished in a group of children. Consequently, Prometheus at the early age must have had a peer companion(s) to invent recursive elements of language and to carry out recursive conversations. A group of children better fits the role of the patriarchs of “unbounded Merge,” than the lone Prometheus.

In some regard, the concept of PFS as uniquely human ability is not new. Uniquely human PFS-like abilities were defined descriptively as “ability to invent fiction”^152^, “episodic future thinking”^153^, “mental scenario building”^154^, “mental storytelling”^155^, “internal mentation”^156^, “mentally playing with ideas”^157^, “creative intelligence”^158^, “prospective memory”^159^, “memory of the future”^160^, “counterfactual thinking”^161^, “integration of multiple relations between mental representations”^12^, “the ability to form nested scenarios”^162^, “an inner theatre of the mind that allows us to envision and mentally manipulate many possible situations and anticipate different outcomes”^162^, “mental exercises that require tracking and integration of what, in the subject’s mind, are temporally separate items of information”^15^; and many more. PFS may be congruent to the “the faculty of language in the narrow sense (FLN)” defined by Fitch, Hauser, & Chomsky (^163^): “Ultimately, we think it is likely that some bona fide components of FLN—mechanisms that are uniquely human and unique to language—will be isolated and will withstand concerted attempts to reject them by empirical research. An understanding of such mechanisms from the genetic, neural and developmental perspectives will illuminate our understanding of our own species…. it seems clear that the current utility of recursive mental operations is not limited to communication. … recursive thought would appear to be quite useful in such functions as planning, problem solving, or social cognition.” PFS was probably the process that Ian Tattersall had in mind when he was talking of the uniquely human “capacity for symbolic thought“:”… If there is one single thing that distinguishes humans from other life-forms, living or extinct, it is the capacity for symbolic thought: the ability to generate complex mental symbols and to manipulate them into new combinations. This is the very foundation of imagination and creativity: of the unique ability of humans to create a world in the mind…”^76^.

Descriptive definitions of PFS-like abilities as uniquely human are not new. The new conjecture proposed in this manuscript is the result of the exact neurobiological definition of PFS. Traditionally, PFS ability is rolled into more general abilities such as executive function, cognition, fluid intelligence, and working memory. None of those traits have a strong critical period since they can be improved well into adulthood^164^. Only by defining PFS as a separate neurological mechanism, were we able to discover the strong critical period for PFS acquisition: individuals who have not acquired PFS in early childhood cannot develop PFS later in life despite years of therapy^4^. Other neurological conditions with a strong critical period include monocular deprivation^63^, filial imprinting in birds^64^ and monaural occlusion^65^.

The entire proposal completely depends on this understanding of the strong critical period for PFS acquisition. (Note that this critical period is different from other language-related critical periods, such as phoneme tuning^56,57^, grammar processing^58^, articulation control^59^, and vocabulary acquisition^60^ that all can be significantly improved by training at any age^61,62^ and, therefore, are weak critical periods.) Less specific, more ambiguous definitions of PFS-like abilities water down its strong critical period and undercut the analysis of language evolution. For example, theory-of-mind (ToM) is often included into PFS-like abilities. Similar to PFS, ToM acquisition has a critical period: deaf children who acquire formal sign language early, are significantly better at reasoning about mental states than language-delayed deaf children^165,166^. ToM, however, improves at any age when individuals learn mental state vocabulary — particularly linguistic forms for verbs such as ‘think’ and ‘know’^166^. Therefore, ToM has a weak critical period and shall not be merged with PFS that has a strong critical period.

Mental rotation and integration of modifiers are often also defined together with PFS since all three active imagination processes are controlled by the LPFC. Similar to PFS, acquisition of mental rotation and integration of modifiers have critical periods^23,26^. However both, mental rotation and integration of modifiers, can be acquired in adulthood following language therapy and therefore have weak critical periods^28,30^. Accordingly, for the purposes of language acquisition, PFS must be considered separately from mental rotation and integration of modifiers.

Similar to other traits with strong critical periods – monocular deprivation, filial imprinting in birds, and monaural occlusion – PFS cannot be acquired in adulthood. Its neural infrastructure has to be laid down in early childhood. Perhaps this neural infrastructure is related to cortical functional specialization established through competition mechanisms similar to that of monocular deprivation^167,168^ and development of long frontoposterior fibers, such as arcuate fasciculus and superior longitudinal fasciculus^169^. The exact mechanism of the strong critical period for PFS acquisition remains to be determined.

### History of language acquisition

The Romulus and Remus hypothesis suggests that the first phase of articulate speech acquisition from around 2 million to 600,000 years ago has to be explained separately from the second evolutionarily fast phase of recursive language acquisition 70,000 years ago (Box 3). Articulate speech relies on multiple neurologically distinct mechanisms each of which is the result of complicated evolution and many genetic mutations. Compared to chimpanzees, modern humans improved neurological control of the diaphragm and the tongue, musculature of the mouth and lips, position and control of the vocal cords, hearing frequency range, neocortical processing of auditory stream, and many other abilities. This piecemeal improvement of articulate speech could not have been fast and probably has taken millions of years.

In some sense, the near-modern speech apparatus circa 600,000 years ago can be viewed as a preadaptation for recursive language. A classic example of pre-adaptation is bird feathers that initially may have evolved for temperature regulation, but later were adapted for flight. The speech apparatus 600,0 years ago served for the purposes of communication, but not for the purposes of unbounded contemplation. Today, once PFS is acquired in childhood, we are able to use the vocabulary both for communication and unbounded contemplations, just like modern birds use feathers for both temperature regulation and flight.

### Neanderthal speech, culture, and hunting styles

We concur with Dediu and Levinson who “attribute to Neanderthals modern speech, double-articulation (separated phonology and lexicon), some systematic means of word combination (syntax), a correlated mapping to meaning, and usage principles (pragmatics).”^1^ However, we refine the description of Neanderthal communication system further. There is no archeological evidence that Neanderthals possessed the PFS ability. Without PFS, their communication system could include nouns, verbs, adjectives, numbers, color and size modifiers, but was void of spatial prepositions, verb tenses, nesting, and other recursive elements. Neanderthal communication system was similar to the communication system used by contemporary individuals who have fluid speech combined with PFS disability: individuals with prefrontal aphasia^15,18^, specific brain damage^12,170–173^, late *first-language* learners^23–26^, and verbal low-functioning individuals^19,174^.

It is likely that further parallels could be drawn between individuals with PFS disability and Neanderthals. The contemporary individuals with PFS disability can be social, compassionate, have normal attention and impulse control, artistic talents, and excellent memory. Similarly, Neanderthals and other pre-PFS hominins could have been social and compassionate. They could have taken care of their sick and diseased^175^. Notably, while empathic contemplations in modern humans are often associated with production of novel mental images, the PFS ability is not necessary for empathy.

Likewise, Neanderthals and other pre-PFS hominins could have been able to understand the concept of symbol. The symbolic use of objects can be associated with PFS in modern individuals, but PFS is not necessary for using an object as a symbol. Symbolic use of red ochre, production of perforated shells^68^, drawing lines and hand stencils do not require PFS. Hybrid art such as lion-man^75^, on the other hand, unequivocally requires PFS^9^ and must have been beyond Neanderthal capabilities.

Both early modern humans and Neanderthals were using animal hides for warmth, but humans could have been stitching those hides into well-fitted clothes with the use of sophisticated bone needles. Stitching clothing definitely relies on PFS since to cut and stitch an animal hide into a well-fitting garment, one needs first to mentally simulate the process, i.e. imagine how the parts can be combined into a finished product that fits the body. Without PFS, Neanderthals must have been simply wrapping the hides around their bodies like a poncho.

The biggest difference in behavior between Neanderthals and modern humans must have been in their hunting styles. As discussed above, building an animal trap is impossible without PFS. Both Neanderthals and modern humans hunted mammoths, but lacking an ability to invent traps, Neanderthals must have tried to puncture animals with as many spears as possible. This style of attack implies close contact with an animal and must have led to a high frequency of bone injury among hunters. Modern humans, on the other hand, had the capacity to chase a mammoth into a pitfall trap where an immobilized weakened animal could have been easily killed. Comparison of archeological remains between early modern hominines and Neanderthals is therefore expected to show disproportionally larger number of broken bones in Neanderthals that must have resulted from this close contacts with large animals^76,176,177^.

### Testable predictions

An important component of a theory is that it should be testable. A theory must make predictions that were not used in the construction of the theory initially but are now available for inspection. If the predictions are borne out, the theory would be strengthened. If not, then the original theory ought to be modified or abandoned. Here we make several predictions derived from the Romulus and Remus hypothesis.

1. Archeological evidence. The Romulus and Remus hypothesis can be disproved by an archeological finding unambiguously demonstrating PFS ability in hominins significantly earlier than 70,000 years ago.
2. Teaching recursive language to an animal. The Romulus and Remus hypothesis predicts that humans are unique in their genetic predisposition to ontogenetic acquisition of PFS. Thus, the hypothesis can be disproved by demonstrating that other living primates are capable of acquiring PFS.
3. Shortened PFC maturation in humans. If *in vivo* biomarkers for PFC maturation rate can be established, then duration of PFC plasticity could be correlated to PFS ability. Individuals with increased rate of the PFC maturation and decreased duration of PFC plasticity are expected to exhibit lower PFS ability. From a theoretical point of view such individuals may significantly benefit from an early intensive language therapy.
4. Effect of passive entertainment on children. Lack of dialogs with the use of recursive language during the critical period is predicted to negatively affect the PFS ability. Passive watching of TV and other videos can significantly reduce time available for dialogs and therefore predicted to result in decreased PFS ability.
5. Children with language delay. Children taking no interest in external and internal language can miss the critical period for PFS acquisition. Child’s non-recursive vocalizations alone cannot inform on PFS acquisition. In these children it is important to assess the PFS function directly^19,178^ and administer intensive language therapy as soon as possible. We are currently conducting an observational trial of the effect of parent-administered PFS exercises in children diagnosed with ASD^179^. We predict that PFS exercises will significantly improve children learning, particularly in the domain of language comprehension.
6. **An artificial extension of the period of plasticity of the PFC in animals**. Recent insights into genetics of the “PFC delay” mutation identified several possible genetic targets enabling this function. Liu X. and colleagues identified four transcription factors that could play a role in regulating the timing of the development of the prefrontal cortex: myocyte enhancer factor 2A (MEF2A), early growth response 1 (EGR1), early growth response 2 (EGR2) and early growth response 3 (EGR3)^145^. MEF2A is predicted to regulate the three EGR genes, as well as several other signal transduction genes^180^. Deleterious mutations of MEF2A were also observed in a significant number of individuals with severe mental retardation^181^. If MEF2A plays a master role in the regulation of the human-specific delay of the PFC maturation, the human version of the gene can be used to extend the period of animals’ PFC plasticity. Chimpanzees with an extended period of PFC plasticity can then be exposed to rigorous cognitive training and recursive language through lexigrams or sign language. A controlled randomized trial comparing chimpanzees with and without the “PFC delay” mutation could demonstrate the influence of the mutation on chimps’ cognitive and language comprehension abilities.
7. **Cloning a Neanderthal child**. In January of 2013, George Church, a Professor of Genetics at Harvard Medical School, said in an interview with the German magazine “Der Spiegel” that it could be possible to clone a Neanderthal baby from ancient DNA if he could find a woman willing to act as a surrogate: “I have already managed to attract enough DNA from fossil bones to reconstruct the DNA of the human species largely extinct. Now I need an adventurous female human.” While currently it is hard to imagine cloning of a Neanderthal for ethical and legal reasons, history teaches us that eventually intellectual curiosity will win over and the Neanderthal will be cloned. How different will it be? George Church suggests, “Neanderthals might think differently than we do. We know that they had a larger cranial size. They could even be more intelligent than us.” Conversely to Church’s conjecture, the Romulus and Remus hypothesis predicts that the Neanderthals were lacking the “PFC delay” mutation and therefore would not be able to acquire PFS and, as a result, will fail to understand spatial prepositions and other sentences describing object combinations, as well as perform below the score of 86 in nonverbal IQ tests. In terms of the neurological difference, Neanderthal brain is expected to feature a smaller superior longitudinal fasciculus as well as a smaller arcuate fasciculus — the frontoposterior tracts that have been shown to be important for PFS^169^.
8. **Human demographic explosion following acquisition of PFS**. The evolutionary mathematical model predicts a demographic explosion following acquisition of PFS 70,000 years ago. First, by changing hunting strategy from persistence hunting to building traps, hominins had the capacity to obtain nearly unlimited quantity of food resulting in increased fertility and decreased mortality. Second, tribe’s losses to predation must have come down since hominins could reduce their exposure themselves to predators during persistence hunting and foraging^182^. Third, the number of wounds received in close combat with large animals had to come down as a result of preferential use of trapping of megafauna. Fourth, PFS must have dramatically increased cohesion between tribe members through religion and recursive language. Fifth, PFS facilitated the process of discovery of new tools, such as spear throwers and bow-and-arrows^183^. The resulting exponential population growth can explain (1) an observed rapid population growth in the ancestral Africa populations around 70,000^184^, as well as (2) unprecedentedly fast diffusion of humans out of Africa 65,000 years ago. The model also predicts increased number of violent deaths as humans with PFS butchered hominins without PFS – the event that could be confirmed by future archeological digs.
9. **Animal traps can appear in archeological record after acquisition of PFS**. It may be possible to identify archeological artifacts of animal traps after 70,000 years ago; perhaps even pit traps characterized by large quantity of animal dung and bones can be identified.
10. **Morphologically-modern versus imagination-modern *Homo sapiens***. Many researchers consider fossils from Morocco dated to 300,000 years ago to be the oldest known examples of the *Homo sapiens* lineage^185^. The discrepancy in the appearance of morphologically-modern and behaviorally-modern *Homo sapiens* is an unsolved puzzle. The Romulus and Remus hypothesis suggest that despite morphological similarity, *Homo sapiens’* imagination was very different from modern imagination until PFS acquisition 70,000 years ago.

## Conclusions

We suggest that the “Upper Paleolithic Revolution” is explained by simultaneous acquisition of recursive language and prefrontal synthesis (PFS), enabled by a genetic mutation that extended the critical period by slowing down the prefrontal cortex (PFC) maturation. Composite figurative art, hybrid sculptures, adorned burials, proliferation of new types of tools, fast diffusion out of Africa into four continents and demise of the Pleistocene megafauna are all logical consequences of the acquisition of the PFS ability. This event completely separated the pre-PFS hominins, who had a non-recursive communication system, from the morphologically similar but behaviorally different breed of hominins who possessed recursive language and relied on mental simulation to plan war and hunting activities. The acquisition of PFS resulted in what was now in essence a behaviorally new species: the first *behaviorally modern Homo sapiens.* The newly acquired power for fast juxtaposition mental objects dramatically facilitated mental prototyping and led to a fast acceleration of technological progress. Humans acquired an ability to trap large animals and therefore gained a major nutritional advantage. As population exploded, humans quickly diffused out of Africa and settled the most habitable areas of the planet. These humans coming out of Africa some 65,000 years ago were very much like modern humans since they possessed both components of full language: the culturally transmitted recursive language along with the innate predisposition towards PFS enabled by the “PFC delay” mutation. Armed with the unprecedented ability to mentally simulate any plan and equally unprecedented ability to communicate it to their companions, humans were poised to quickly become the dominant species.

### Box 1. Who were the children who invented recursive elements of language and acquired PFS?

Who were the children who first invented recursive elements and acquired PFS? Three conjectures are fairly certain: (1) they must have lived together for several years, so that they could often talk to each other, (2) they must have had acquired the novel “PFC delay” trait, and (3) one of them must have survived to adulthood to have children of his own and to teach them recursive language. Other inferences are speculative: These children-inventors were unusually slow compared to their peers. Slow children do best in highly supportive environment, so maybe they were children of a chieftain. They spent a lot of time together and created their own vocabulary, so maybe they were closely spaced siblings or even twins; twin bonding and their propensity for cryptophasia are well documented^138^. The “PFC delay” trait could have been caused by a *de novo* germline dominant mutation or combination of their parents’ recessive mutations.

We attempted to reject the least probable scenario of monozygotic twins sharing the dominant *de novo* “PFC delay” mutation. The following calculations show that the low monozygotic twins birth rate and the low probability of the specific “PFC delay”-causing mutation are not enough to reject the monozygotic twins’ hypothesis. Monozygotic twins birth rate probability is about 0.003 and is uniformly distributed in all populations around the world^186^. At a minimum, one twin had to survive to the age of 4 (probability=0.58) and another twin had to survive to the age of 20 to have children of his own and to teach them recursive language (probability=0.22). Thus, the surviving twins’ probability = 3.8×10^-4. Assuming 1000 hominin tribes and 16 births per tribe per year, six monozygotic twins are expected to survive yearly.

Probability of any birth with a *de novo* dominant mutation affecting any transcription factor can be calculated as follows. Humans have approximately 100 new mutations per birth^187^. Conservatively assuming that only 10 out of 3×10^9 base pairs result in the change of function of the “PFC delay”-controlling transcription factor, the probability of the “PFC delay” mutation = 100 x 10 / 3×10^9 = 3.3×10^-7. Again, conservatively assuming 1000 hominin tribes^188^ and 16 births per tribe per year, in 500,000 years between acquisition of the modern speech apparatus and PFS acquisition, the “PFC delay” mutation occurred in at least 2640 births. Multiplying this number by the probability of the surviving twins, we calculate that at least one set of surviving twins was born with the *de novo* dominant “PFC delay” mutation in 500,000 years. Thus, it is possible that Romulus and Remus, the inventors of recursive language, were monozygotic twins (Box 2).

### Box 2: A legend of Romulus and Remus written by Matthew Arnold

Romulus and Remus were twin brothers born to Kital, the chief of a tribe of early hominins. After multiple attempts to produce a male heir, he finally got what he wanted, but soon discovered that his children were ‘different.’ When they were born, Romulus and Remus seemed like every other child physically, but odd enough they seemed to lack common sense. They would go near dangerous tar pits, animal dens, and rivers. At the age of 4, they even wandered out into the forest and got lost, something even younger children would not do. Most kids their age already had basic knowledge of their role in society and a simple understanding of what to do and what not to do out in the wild, but Romulus and Remus had no such understanding. While, most children would understand not to go near a rushing river to gather water, the twin brothers almost drowned because they lacked the basic knowledge not to do so. Due to his children’s intellectual inadequacies Kital was extremely displeased with his sons. He wanted great warriors that would grow up to lead his tribe, but unfortunately, he was growing older and after so many failed attempts to produce a male heir, he had no choice but to try and raise his children the best he could. However, as the head of his tribe he lacked the time to constantly look after his sons, so he got the tribe’s medicine man to do so for him in the hope some of his wisdom would rub off on the young boys.

A few years later, while under the care of the medicine man the brothers began to speak the primitive language of their fellow tribe members and slowly picked up on the social cues and common knowledge of others. However, the brothers went a step beyond, they started to add spatial prepositions to their language, something never done before. Even the medicine man did not know what to make of the addition of these new words to the children’s language, and since he could not understand what they meant, he just assumed they were fooling around. The medicine man would also notice strange drawings around the cave where the boys spent most of their time. There were markings carved into the walls of the cave, some of which resembled animals. A normal tribe member would dismiss these drawings as pointless but not the medicine man. He thought these drawings were incredible and illustrated something special about the young boys. Just like the chief, the medicine man did not have much confidence in Romulus and Remus, but their newfound ability changed his mind, and it was not long until the medicine man deemed the boys ready to embark on a hunt on their own, a rite of passage so to speak.

The medicine man and Kital brought them to the hunting grounds and gave them each a spear. It was now time for them to prove themselves worthy successors or die trying. Even though he was the one that agreed to this test, Kital was still terrified that his sons would not be able to catch anything or be killed by wild beasts and his last chance to have an heir would vanish, but what happened astounded him. When he told them that they would have to catch a buffalo on their own he saw them discuss a plan to catch the animal, but oddly enough, despite speaking the same language, Kital did not recognize many of the words they spoke. He could understand words like “buffalo,” “run,” and “rocks,” but he could not understand the relationship between the words. The boys then went to a small path with steep cliffs overlooking both sides and drew a circle in the dirt and dug a hole where the circle was and covered it with leaves and branches so that it was indistinguishable from the surrounding area. Kital was confused by his sons’ behavior, because normally hunters would utilize persistence hunting, which is when hunters would chase animals until the creature would die of exhaustion. After creating the hole, the brothers conversed again, and ran to find buffalo. Around half an hour later, they chased a group of them back to the corridor where they had dug the hole, and as the buffalo passed over the leaves and sticks, they fell down into the pit. Immediately after, the two brothers speared the trapped animals and pulled them out. The brothers had caught not just one wild buffalo, but many. Simply catching one buffalo would have taken a normal hunter, hours to accomplish by use of persistence hunting. It was at this moment that Kital knew he had somehow succeeded in producing successful male heirs.

As years went by, the brothers improved the language they spoke, introducing more spatial prepositions and eventually developing a complete recursive language. Even though their father could not fully understand them, the brothers could work and communicate together to produce miraculous results. This led them to gain the respect of their fellow tribe members as well as their father. Together, Romulus and Remus would end up leading the tribe and conquer all the neighboring tribes with their enhanced intelligence and ability to formulate complex military tactics. Eventually, when Romulus and Remus had children, they found they were just like them when they were young, and they were able to teach them the language they had created. They would take care of their children for a longer time, but after several years, they were able to use the more complex language that their parents had created. These two brothers unknowingly started a pattern that would continue for tens of thousands of years and lead to the modern humans of today.

### Box 3: History of verbal communication

The Romulus and Remus hypothesis divides the history of language acquisition into two phases: the nearly 2-miilion-year-long period of gradual growth of vocabulary made possible by piecemeal improvements of the vocal apparatus, as described by Dediu and Levinson^1^, and a fast several-generations-long, conversion to modern recursive language 70,000 years ago. This info box charts a possible time course of polygenic vocabulary acquisition in the genus *Homo* from 2 million years to about 600,000 years ago and describes possible evolutionary forces that shaped the modern speech apparatus.

*Homo erectus* was an adventurous species with the body built for long distance running^189^. In fact, *Homo erectus* was moving so much that the species diffused out of Africa and settled most of Europe and Asia starting around 1.8 million years ago^190–192^. Any modern camper traveling in a group would appreciate the value of effective communication. In each new place, the group needs to locate a protective shelter, edible food, a source of clean water and carry out a myriad of other location-specific projects. Any improvement in vocabulary would have allowed *Homo erectus* chieftain to delegate jobs much more efficiently. A leader could purposefully select someone more suitable for the job: he could select John, because John is stronger, or Peter because Peter is taller, or Steve who can climb trees faster. A leader who found a cave with a prohibitively small opening would ask John to come with him and help clear the large boulders from the cave’s entrance, but ask Steve to climb a tree if he has found a beehive full of honey. Such arrangement could significantly benefit from personal names. We envision that some mutation improved neurological control of the diaphragm or tongue, musculature of the mouth or lips giving that individual slightly better mechanical ability to articulate sounds and enabled him to increase the number of distinct words from chimpanzee’s 20 to 100^110–113^ to 150. Hominins lived in groups of 50 to 150 individuals and 150 words could have been used to assign names for everyone in a group (after all, even dolphins have life-long given names called signature whistles^193,194^). Even if no one else in the group but the leader was able to call each person by name, the leader and the group as a whole would have gained an advantage of an increased cooperation and therefore better nutrition. As an alpha male, the leader would have high number of children and thus his “improved vocal apparatus” mutation would have had fixed in a population.

After many generations another leader could get a different mutation that further improved his vocal apparatus and doubled his vocabulary to 300 words. The articulate leader could have been using extra words as nouns to further facilitate job assignment without the need to point to each object: two-word sentences could communicate job assignment: “John flint,” meaning that John is expected to collect flint stones; “Peter sticks,” meaning that Peter is expected to find sticks; “Patrick tubers,” meaning that Patrick is expected to dig tuber; and so on. The leader could also instruct the selected workers in what to take with them: handaxes for cutting trees, spears for hunting, or a sack for carrying throwing stones back to the shelter.

Thousands of years later another mutation may have extended vocabulary of the leader to 600 distinct words and enabled the leader to name more objects, tools, and actions, further improving his ability to efficiently organize the tribe’s productive activities. This leader could have now used more complex sentences: “John, come here;” “Peter, bring the handaxe;” and “Patrick, collect stones.” Different types of edible plants and prey animals could have been assigned their own names and different jobs could have been assigned action words. Various geographical locations, including rivers, mountains, caves, and maybe even individual trees could be named helping adventurous *Homo erectus* to orient and to describe directions to other tribesmen.

Thus, we envision development of vocal apparatus as a series of beneficial mutations slowly improving control of diaphragm, lips, tongue, chicks, vocal cords larynx position in the trachea, and possibly hundreds of other related mutations over many millennia. When articulate speech mutations originate in a leader, they result in immediate improvement of communication (albeit one-way communication) between tribe members and consequent increase in productivity (aren’t our leaders still more articulate than an average person?). Since leaders also had higher chances to procreate, their “improved vocal apparatus” was slowly fixed in a population over many generations.

Critically, such a communication system with many names, nouns and verbs, while significantly benefiting traveling parties of *Homo erectus,* does not rely on PFS. In fact, chimpanzees, dogs and some other animals have been trained to follow hundreds of commands, such as “bring the ball/newspaper/slippers,” “leak the hand/floor/bowl,” that rely on memory of nouns and verbs, but do not require purposeful combination of objects from memory into novel mental images via PFS.

The next hypothesized step in the evolution of language is beyond the ability of any living animal, but still does not rely on the PFS ability. It involves acquisition of the type of imagination called integration of modifiers^2^. Anyone reading this manuscript can obviously imagine their room bigger or smaller, painted blue, red, or yellow and follow the instruction to find a *long red* straw placed among several decoy objects including other *red* shapes (Lego pieces, small *red* animals) and *long/short* straws of other colors. Integration of modifiers is a type of active imagination relying on the LPFC ability to change the size and color of objects in front of the mind’s eye. Integration of modifiers was never demonstrated in any animal and is missing from about 7% of modern humans (7% is the number of humans not capable of answering questions requiring integration of size and objects, or color and objects when tested with matrix reasoning tasks of typical nonverbal IQ test, see^5^.

Acquisition of integration of modifiers ability could have been also influenced by *Homo erectus* mobile lifestyle. Whether foraged or hunted, *Homo erectus* would have exhausted the land around their shelter within a few months and therefore must have traveled regularly from one place to another. For a group of 50-150 hominins, wandering around without knowing the final destination is highly inefficient. Exacerbating the problem would be the fact that most women in the group would have been either nursing or pregnant. Looking for a new shelter in the company of pregnant or nursing women and small children would have been highly dangerous. It is safer and more efficient to decide on an ideal location first, and then gather everybody together to make a quick move. Therefore, it is likely that able-bodied scouts were sent out to look for the next fertile area with a nearby shelter, while the rest of the group stayed behind in safety. Adjectives and integration of modifiers ability would have allowed scouts returning from their trip to compare their observations and decide not only which shelter was better (e.g., a bigger cave, or one with a nearby water source), but also which shelter was best positioned as far as availability of prey animals and edible plants (grasses, herbs, seeds, roots, rhizomes, or tubers). Such discussions and comparisons still do not require PFS. For example, scouts could have measured the size of their caves using their own strides (the size of which is comparable in similarly-built individuals) and then communicate this information to the leader to help choose the largest cave. A tribe leader capable of integration of modifiers would be able to select a better pasture and a better shelter and therefore would have been able to father greater number of healthy children passing down his genes for improved language.

We conclude that *Homo erectus* lifestyle was conducive to improvements of vocal apparatus. *Homo erectus* could have used a communication system with thousands of words without PFS. It is possible that *Homo erectus* may have even had more words than modern humans since without recursion, they needed more words to communicate the same number of concepts.

## Acknowledgments

We wish to thank Dr. P. Ilyinskii, Dr. K. Khrapko, and Dr. D. Gamarnik for productive discussion and Dr. P. Ilyinskii for scrupulous editing of this manuscript.

